# A Focused Small-Molecule Screen Identifies PHA-680626 as an Amphosteric Inhibitor Disrupting the Interaction between Aurora-A and N-Myc

**DOI:** 10.1101/2021.03.10.433854

**Authors:** Fani Souvalidou, Dalila Boi, Roberta Montanari, Federica Polverino, Grazia Marini, Davide Capelli, Giorgio Pochetti, Roberto Contestabile, Daniela Trisciuoglio, Angela Tramonti, Patrizia Carpinelli, Camilla Ascanelli, Catherine Lindon, Alessandro De Leo, Roberto Di Santo, Roberta Costi, Giulia Guarguaglini, Alessandro Paiardini

## Abstract

Neuroblastoma is a severe childhood disease, accounting for ~10% of all infant cancers. The amplification of the MYCN gene, coding for the N-Myc transcriptional factor, is an essential marker correlated with tumor progression and poor prognosis. In neuroblastoma cells, the mitotic kinase Aurora-A (AURKA), also frequently overexpressed in cancer, prevents N-Myc degradation, by directly binding to a highly conserved N-Myc region, i.e. Myc Box I. As a result, elevated levels of N-Myc, which are required for the growth of MYCN amplified cells, are observed. During the last years, it has been demonstrated that the ATP competitive inhibitors of AURKA CD532, MLN8054 and Alisertib also cause essential conformational changes in the structure of the activation loop of the kinase that prevent N-Myc binding, thus impairing the formation of the AURKA/N-Myc complex. In this study, starting from a screening of crystal structures of AURKA in complex with known inhibitors, we identified additional compounds affecting the conformation of the kinase activation loop. We assessed the ability of such compounds to disrupt the interaction between AURKA and N-Myc *in vitro*, using Surface Plasmon Resonance competition assays, and in tumor cell lines overexpressing MYCN, by performing Proximity Ligation Assays. Finally, their effects on N-Myc cellular levels and cell viability were investigated. Our results, identifying PHA-680626 as an amphosteric inhibitor both *in vitro* and MYCN overexpressing cell lines, expand the repertoire of known conformational disrupting inhibitors of the AURKA/N-Myc complex, and confirm that altering the conformation of the activation loop of AURKA with a small molecule is an effective strategy to destabilize the AURKA/N-Myc interaction in neuroblastoma cancer cells.

## INTRODUCTION

AURKA, a member of the Aurora Kinases family, is a Ser/Thr kinase involved in the progression of mitosis. In interphase it associates with centrosomes and controls their maturation in the G2 stage of the cell cycle, while in mitotic cells it localizes at spindle poles, where it plays a primary role in establishing spindle bipolarity and regulating microtubule nucleation and dynamic properties (Joukov & De Nicolo, 2018; Magnaghi-Jaulin et al., 2019). AURKA is widely overexpressed in a variety of solid tumors (Bischoff et al., 1998; Sen et al., 2002; Du et al., 2021). In particular, in neuroblastoma, a severe childhood cancer that arises from highly proliferative migratory cells of the neural crest (Grimmer & Weiss, 2006), AURKA is not only highly expressed relative to normal tissues (Zhou et al., 1998), but also displays another critical function, by binding to and stabilizing the oncoprotein N-Myc (Otto et al., 2009).

N-Myc belongs to the MYC family together with C-Myc and L-Myc, a very potent network of transcriptional factors, regulating the expression of a massive group of genes implicated in cell cycle progression, protein translation and metabolism (Albihn et al., 2010; Grandori et al., 2000). Such transcriptional factors are usually found upregulated in various cancers in humans (Dang, 2012). The structural organization of Myc proteins shares conserved motifs called “Myc Boxes” (MB), which serve as a docking site for protein-protein interactions. The turnover of Myc proteins is regulated by the phosphorylation of MBI (Myc Box I), which directs the protein towards ubiquitination and proteolysis (Welcker et al., 2004). N-Myc is primarily phosphorylated on Ser62 by the Cdk1/Cyclin-B complex, and then also on Thr58 by the Gsk3 kinase (Yada et al., 2004; Sjostrom et al., 2005; Chesler et al., 2006). Dephosphorylation of Ser62 by the PP2A phosphatase recruits the activity of E3 ubiquitin ligase SCFFbxW7 (Welcker et al., 2004; Adhikary & Eilers, 2005). The physical interaction of N-Myc with AURKA, as demonstrated by *in vitro* and crystallographic studies, does not allow the ubiquitin ligase to intervene, thus preventing N-Myc degradation (Richards et al., 2016).

During the years, serious efforts have been taken to exploit specific methods for targeting AURKA in cancer, mainly focused on the design of various inhibitors that are under investigation in clinical trials (Malumbres & Ignacio, 2014; Levinson, 2018). Apparently, an important subclass of these inhibitors, hailed as “conformation disrupting” (CD) or “amphosteric”, is also able to effectively disrupt the protein-protein interaction of the kinase with N-Myc, promoting the degradation of N-Myc itself (Gustafson et al., 2014; Meyerowitz et al., 2015; Brockmann et al., 2015). The binding of such inhibitors induces a series of unique interactions that produce a conformational change on the structure of the kinase domain of AURKA, distinct from any known physiological state (Meyerowitz et al., 2015). Notable examples come from CD532, MLN8054 or MLN8237 (Alisertib), diaminopyrimidine derived compounds that disrupt the interaction between AURKA and N-Myc, due to a significant shift of the N-terminal domain relative to the C-terminal one, and most importantly to the repositioning of the kinase activation loop (the “A-loop”, comprising residues 276-291 of human AURKA) in an inactive (hereinafter “closed”) conformation, which prevents N-Myc binding (Gustafson et al., 2014; Richards et al., 2016).

Beside their classification as “orthosteric” (inhibiting kinase activity), “allosteric” (disrupting protein-protein interactions) or “amphosteric” (both), in general kinase inhibitors are classified as “type I” or “type II”, according to their preference for a “DFG-out” or “DFG-in” target state, which refers in turn to the orientation of a catalytically important motif of three residues, i.e. Asp(D)-Phe(F)-Gly(G), found at the N-terminus of the kinase A-loop (Schindler et al., 2000; Pargellis et al., 2002; Wan et al., 2004). In the past, the closed state of the A-loop was invariably linked to the “DFG-out”-associated, type II inhibitors. However, recent studies have demonstrated that this is not a strict rule. For example, CD532, which is known to confer the closure of the A-loop, keeps the motif in state “DFG-in”. Instead, a highly potent aminopyrimidinyl quinazoline AURKA type II inhibitor is not able to stabilize the A-loop in a closed state (Heron et al., 2006; PDB Code: 2C6E), as well as the potent MK8745 type-II compound, which does not act as a CD inhibitor (Lake et al., 2018).

Here we investigated, by computational and experimental means, which compounds already known to target AURKA at the ATP binding site (i.e., “orthosteric” compounds), regardless of their “type I” or “type II” nature, are able to act as conformation disrupting inhibitors, and destabilize the complex with N-Myc. An *in-silico* data mining in PDB of all known AURKA inhibitors was conducted, to investigate which molecules are capable of opening the “angle” of the kinase in a similar way to other CD inhibitors, as well as to close the A-loop. A series of identified compounds potentially acting as CD inhibitors were firstly confirmed with Surface Plasmon Resonance (SPR) and kinase activity assays to bind to AURKA at the active site and to inhibit its enzymatic activity. Using SPR competition assays with N-Myc, we demonstrate that some of those compounds so far known only as inhibitors of the kinase activity of AURKA, are indeed able to disrupt also the Aurora-A/N-Myc complex *in vitro*. Finally, those compounds were also tested in MYCN overexpressing cell lines, for their ability to disrupt the complex. The most important candidates, together with CD532, MLN8054 and Alisertib, represent new potential pharmacological tools to disrupt the “unholy matrimony” between two proteins that are mainly responsible for the severity and poor outcome of neuroblastoma.

## RESULTS

### PHA-680626 and RPM1722 are predicted to prevent N-Myc binding to AURKA

By screening the whole Protein Data Bank (PDB) subset of AURKA in complex with small molecule compounds, we initially superposed each structure with apo-AURKA (PDB: 4J8N) and with the AURKA/N-Myc complex (PDB: 5G1X) (Richards et al., 2016), and identified several entries in which both of the following conditions were satisfied: 1) AURKA opens its N-terminal domain relative to the C-terminal one (the opening was assessed by measuring the angle between the α-carbons of Val324, Glu308, and Ala172, as already described in Meyerowitz et al., 2015); 2) the A-loop of AURKA is in a state which is incompatible with N-Myc binding, due to the presence of active site inhibitors. Among these, we selected CD532 and MLN8054, which were already shown to decrease the amount of N-Myc in neuroblastoma cell lines (Brockmann et al., 2013; Gustafson et al., 2014) (**Figure 1**). Alisertib (MLN8237), whose structure in complex with AURKA has not been solved yet, was also added for its very high similarity with MLN8054. Moreover, the effects of CD532, MLN8054 and Alisertib have been already attributed to specific conformational changes of the kinase. Two additional compounds, i.e. PHA680626 and RPM1722 (Fancelli et al., 2006; Martin et al., 2012), already known for being orthosteric inhibitors of AURKA, but not investigated, to our knowledge, for their conformation disrupting potential, were selected for further investigation, since the opening of the kinase lobe was particularly high (≥ 85°), and the A-loop was stabilized in a closed conformation (**Figure 1**).

**Figure 1.**
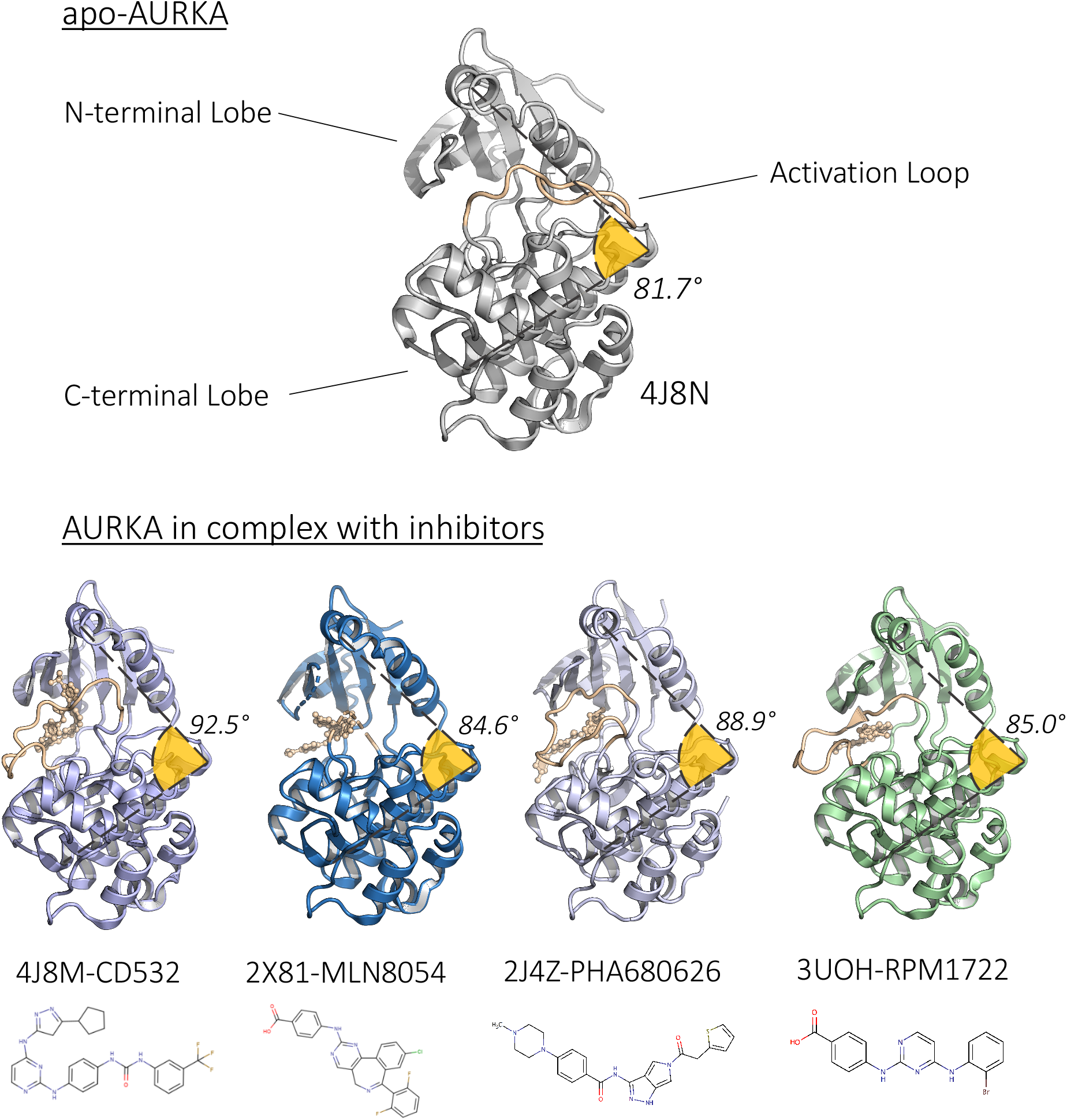
Structures of human AURKA in complex with the investigated inhibitors. The activation loop (AL) is shown in orange. The angle between the N- and C-lobes, measured between α-carbons of V324, E308, and A172, is also shown. The chemical structures of the compounds investigated is also represented.

PHA-680626 derives from the optimization of PHA-680632 (Soncini et al., 2006), a compound based on a 1,4,5,6-tetrahydropyrrolo[3,4-c]pyrazole bi-cycle scaffold, which is known to target the ATP-binding pocket of kinases. The 5-amidothiophene substituent of PHA-680626 is directed away from the glycine-rich loop and packs around Leu263 and Ala273, in proximity of the DFG-motif. A key interaction stabilizing the A-loop in its closed state involves the stacking between the thiophene ring of PHA-680626 and His280 of the A-loop (Fancelli et al., 2006). RPM1722 relies on the bis-anilinopyrimidinic scaffold that is typical of several “DFG-in” inhibitors; however, a bromine substituent on the benzene ring of the scaffold, directed at the DFG flanking residue Ala273, is responsible for induced-dipole forces along alanine side chain, which are reflected by drastic conformational changes of the A-loop. This unique “DFG-out/loop-in” conformation is also stabilized by hydrogen bonding interactions between RPM1722 and residues Lys141 and His280, and by a conformational shift of Trp277, moving from a polar environment to a hydrophobic pocket (Martin et al., 2012).

These observations confirm that other molecules, in addition to the already known amphosteric AURKA inhibitors, induce conformational changes at the kinase lobes and a complete flip of the A-loop, retaining AURKA in a closed conformation that could prevent N-Myc binding.

### PHA-680626 and RPM1722 behave as CD inhibitors *in vitro*

In order to verify that PHA-680626 and RPM1722, and the previously identified inhibitors (CD532, Alisertib, and MLN-8054), were active, *in vitro* kinase activity assays on the purified kinase domain of AURKA were carried out (**Table 1** and **Supplementary Figure 1**). Ten different concentrations of each compound (solubilized in 100% DMSO) were used in the presence of a concentration of ATP equal to the K_M_ reported in the literature (Dodson & Bayliss, 2012)(see Materials and Methods). The measured IC_50_ values were comparable to those found in literature (Martin et al., 2012; Gustafson et al., 2014).

**Table 1.**
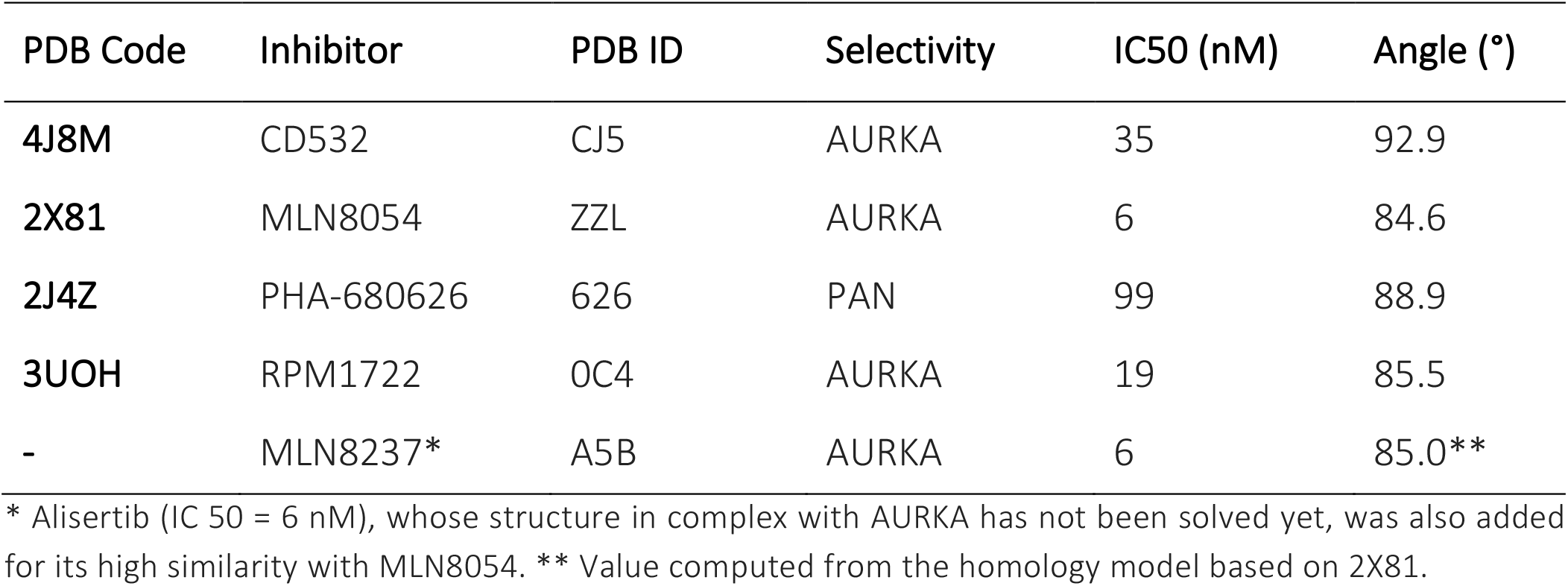
List of AURKA inhibitors investigated in this study*.

In parallel, we also evaluated the interaction between the inhibitors and AURKA by Surface Plasmon Resonance (SPR) binding kinetic assay. AURKA was immobilized onto a PCH sensor chip (see Materials and Methods) and each compound was injected at increasing concentrations in running buffer (10 mM Hepes pH 7.4, 150 mM NaCl, 1% DMSO, 0.005% Tween 20). The interaction between AURKA and the ligands was indicated by the increase of Resonance Units (RUs) compared to the baseline. As the dynamic equilibrium of the A-loop conformation of AURKA influences the binding of small molecule inhibitors, in order to minimize the low affinity binding mode, we carried out the experiment at a fast flow rate (150μl/min) and, again, obtained for all compounds Kd values in agreement with data from literature (**Table 2** and **Figure 2A-E**). An SPR binding kinetic assay was then used to evaluate the binding of the AURKA interacting region (AIR) of N-Myc (N-Myc residues 61–89) (Richards et al., 2016) to AURKA at the same conditions of flow rate and running buffer as for the inhibitors. A Kd value of ~1 μM was obtained (**Figure 2F**). After determining the affinity constants, we performed SPR competition experiments to evaluate the conformational disrupting potency of the inhibitors and their ability to interfere with N-Myc binding to the kinase. In particular, a saturating concentration of each inhibitor (at least 10 times higher than the Kd) was first added to the running buffer to ensure that all the sites of AURKA were filled. Then, N-Myc was injected at different concentrations (125 nM, 250 nM, 500 nM, 1 μM, 2 μM, 4 μM, 8 μM, 16 μM) at a constant flow rate (150 μl/min) of running buffer. As a negative control, we also tested one of the most potent and specific inhibitors of AURKA, MK8745 (an analog of MK-5108 with IC_50_ of 0.6 nM) and ZM447439 (a pan-Aurora family inhibitor, with a measured IC_50_ of ~100 nM) (Georgieva et al., 2010; Chowdhury et al., 2012). Results in **Table 2** indicate that the selected inhibitors almost completely prevent N-Myc from binding to AURKA (Kd_N-Myc_ > 15 μM or not measurable), thus demonstrating their behavior as conformation disrupting compounds, and confirming our initial hypothesis. In agreement with the structural analysis, MK8745 and ZM447439 do not act as CD inhibitors and they do not interfere with the AURKA/ N-Myc interaction (**Table 2** and **Figure 3**).

**Table 2.** SPR analyses of ligands binding. Full kinetic analysis of the AURKA inhibitors.

| Inhibitors | $K_{on}(M^{-1}s^{-1})$ | $K_{off}(s^{-1})$ | $K_d(nM)$ | Res SD | $K_{d-N-Myc}(\mu M)$ |
| --- | --- | --- | --- | --- | --- |
| CD532 | 3.58±0.05e6 | 0.0937±0.0007 | 26.2±0.4 | 0.83 | No binding |
| MLN8054 | 2.02±0.03e7 | 0.0461±0.0004 | 2.28±0.03 | 1.254 | No binding |
| PHA-680626 | 9.15±0.04e5 | 7.83±0.02e-3 | 8.56±0.04 | 3.7 | No binding |
| RPM1722 | 2.61±0.06e6 | 0.013±0.0001 | 5.0±0.2 | 3 | No binding |
| Alisertib | 8.2±0.1e4 | 5.04±0.04e-3 | 60±1.0 | 2.3 | 48 |
| MK8754 | 7.8±0.1e6 | 0.01205±0.00009 | 1.55±0.03 | 1.65 | 0.7 |
| ZM447439 | 9.6±0.06e4 | 9.07±0.04e-3 | 92±0.7 | 1.09 | 4 |

**Figure 2.**
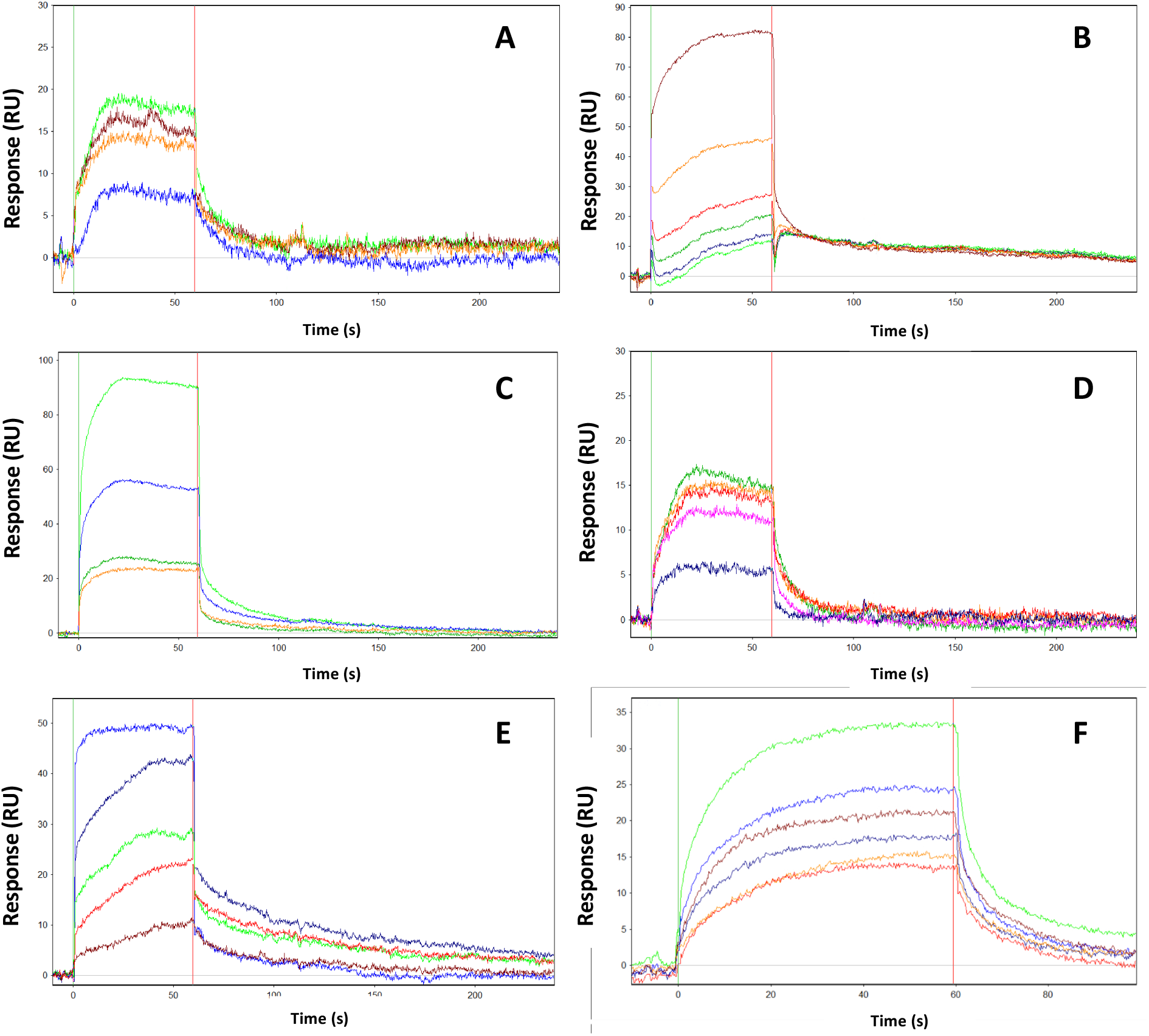
SPR analyses of ligands binding to AURKA. Full kinetic analysis of the AURKA inhibitors CD532 (A), MLN8054 (B), PHA-680626 (C), RPM1722 (D), Alisertib (E), and N-Myc (F). Determined binding parameters are listed in Table 2. Kinetic data for N-Myc: K_on_, 6.3±1e4 M^-1^s^-1^; K_off_, 0.0659±0.0008 s^-1^; Kd, 990 nM; Res SD 2.17. The interaction between AURKA and the ligands is indicated by the increase of Resonance Units (RUs) on the vertical axis compared to the baseline.

**Figure 3.**
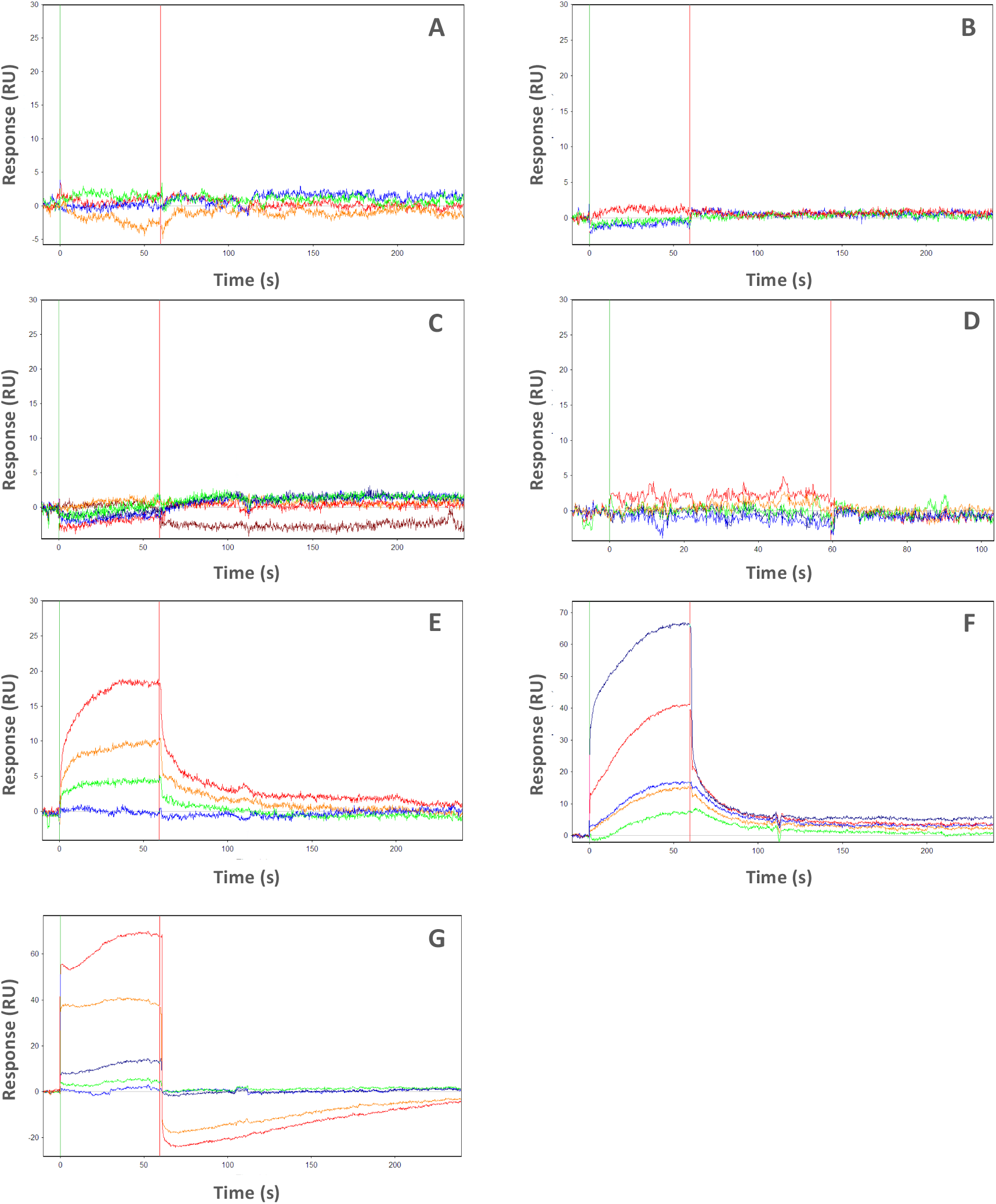
Sensorgrams of competition experiments between the inhibitors CD532 (A), MLN8054 (B), PHA-680626 (C), RPM1722 (D), Alisertib (E), MK8754 (F), ZM447439 (G) and N-Myc. The experiment was carried out by injecting N-Myc at concentrations ranging from 125 nM to 16 μM. N-Myc was previously diluted in running buffer containing a saturating concentration of the inhibitor (at least 10 times higher than its Kd). Apparent Kd values of N-Myc are shown in Table 2. Apparent Kd values for the negative controls MK8754 and ZM447439 are 690 nM and 4 μM, respectively.

### PHA-680626 behaves as a CD inhibitor in MYCN overexpressing cell lines

We then wanted to test whether PHA-680626 and RPM1722 were able to act as CD-inhibitors, and thus to inhibit the N-Myc/AURKA interaction, in cultured cells. Since the interaction is reported as cell cycle-dependent (Buchel et al., 2017) and may vary depending on the levels of the two proteins in cancer cells, we employed a U2OS cell line, that does not express endogenous N-Myc, engineered for inducible doxycycline-dependent expression of mNeon-tagged N-Myc (**Figure 4**). Cultures treated with 1 μM of each compound and with doxycycline were synchronized in the late G2 phase, when AURKA levels are highest, by the RO-3360 Cdk1 inhibitor (protocol in **Figure 4A; Supplementary Figure 3A**). Interestingly, N-Myc failed to accumulate to levels observed in control cultures (DMSO) after treatment with PHA-680626, suggesting an impaired interaction with AURKA that destabilizes it (**Figure 4B**). To directly measure the interaction between AURKA and N-Myc under these conditions, *in situ* Proximity Ligation assay reactions were set up (**Figure 4C** and **Supplementary Figure 3B**), adding the proteasome inhibitor MG-132 in the last 4 hours before fixation to counteract potential destabilization effects. After treatment with small molecule inhibitors *is*PLA signals were counted within cell nuclei and compared to control cells. Results indicate that cells with more numerous *is*PLA interaction spots are less represented in PHA-680626-treated cultures, to a higher extent compared to MLN8054, while RPM1722 is ineffective (**Figure 4C**.). This is accompanied by an increase in the percentage of cells with low *is*PLA signals after PHA-680626 treatment, indicating an impaired AURKA/N-Myc interaction. We then moved to a neuroblastoma MYCN-amplified cell line, i.e., IMR-32. Treatment for 48 hours with 1 μM PHA-680626 decreased N-Myc levels in a comparable manner to MLN8054; no major effect was observed with RPM1722 treatment (**Figure 5A**). By FACS analysis PHA-680626 yielded an increase in the fraction of cells with G2/M DNA content, similarly to known AURKA inhibitors (i.e., MLN8054, **Figure 5B**), while RPM1722 did not induce major cell cycle changes. Interestingly, in contrast to RPM1722, PHA-680626 administration affected IMR-32 cells morphology (**Figure 5C**), with the disappearance of viable substrate-adherent cells and the formation of cellular aggregates that detach from the surface. Consistent with induction of cell death, PHA-680626 and MLN8054 treated cultures displayed a fraction of cells with sub-G1 DNA content (about 40% and 30% respectively, compared to 8% and 15% in DMSO or RPM1722 conditions), as well as a fraction of cleaved PARP-1 in western blot analysis (**Figure 5A** and **5B**).

**Figure 4.**
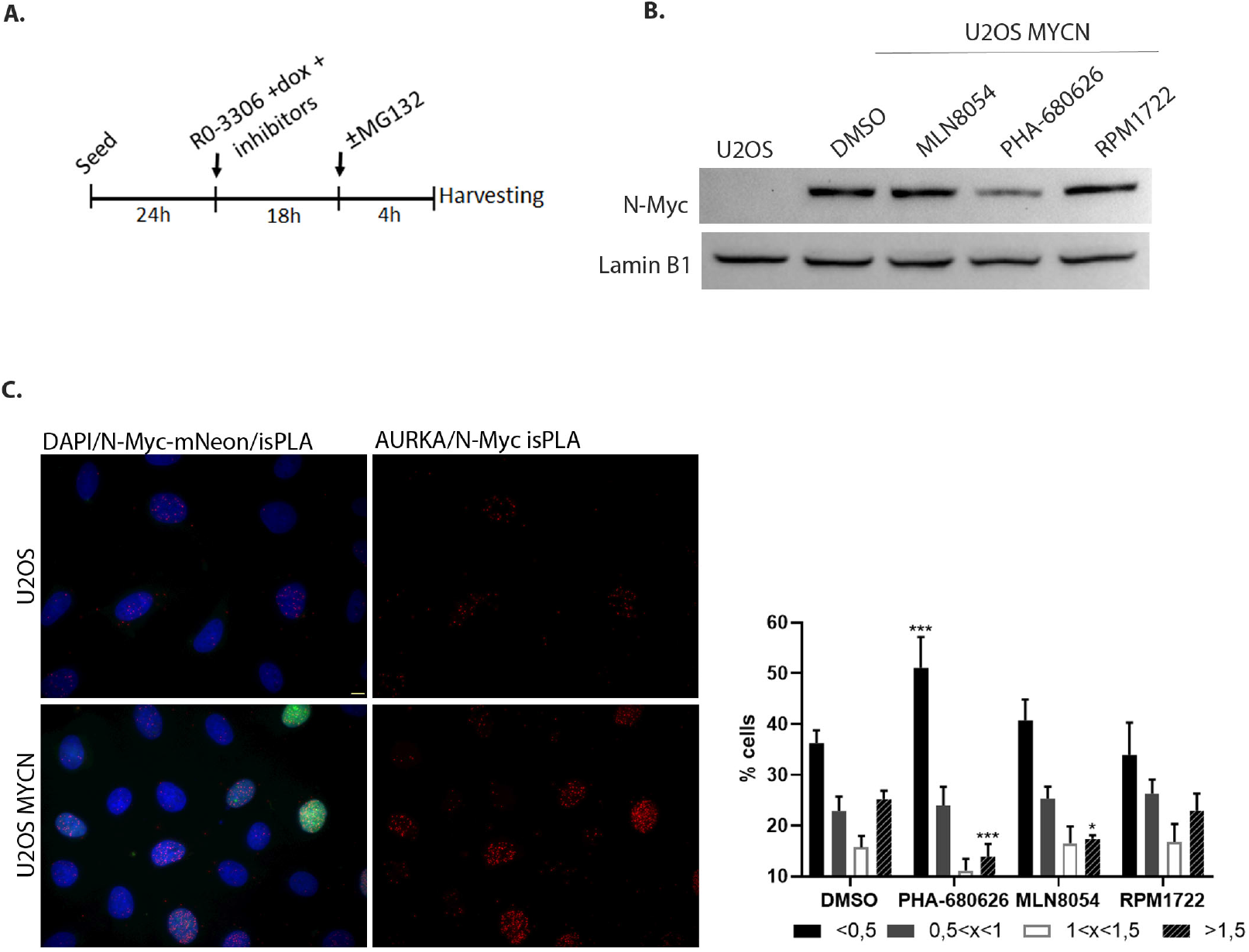
PHA-680626 is able to decrease AURKA/N-Myc interaction. (A) Schematization of the combined synchronization and treatment protocol. (B) Western Blot analysis from U2OS and U2OS MYCN cell lines treated with 1 μM of each compound, or DMSO as control, following the protocol described in (A), without MG132. Lamin B1 was used as loading control. (C) *in situ* Proximity Ligation assays to visualize the AURKA/N-Myc complex formation. The fluorescence panels show examples of the negative (U2OS that do not express N-Myc) and the positive (U2OS MYC dox-induced) reference cultures. The histograms on the right show the distribution (%) of cells in classes, defined by *is*PLA spot values. Values represent isPLA spots per nucleus, normalized to the average value in control cells (at least 1400 measured cells per condition, from 4 independent experiments); mean and standard deviation are shown. Statistical analysis by 2way ANOVA-Tukey’s multiple comparison, *: p<0.05; ***: p<0.001. Scale bar: 10 µm.

**Figure 5.**
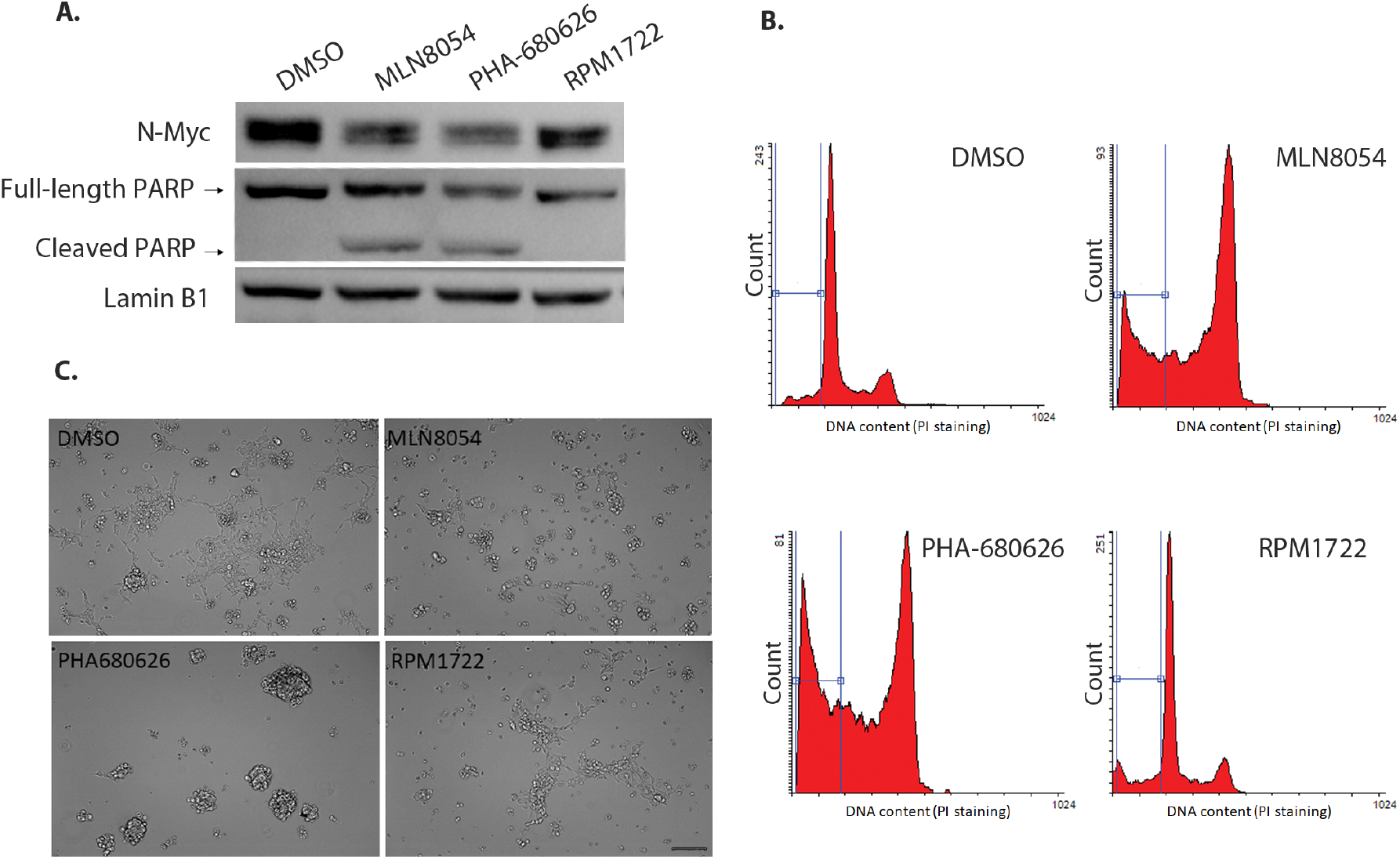
PHA-680626 causes N-Myc protein decrease and cellular stress in neuroblastoma cells. (A) Western Blot analysis from IMR-32 neuroblastoma cell line treated with 1 μM of each inhibitor, or DMSO for control, for 48 h. Both full length and cleaved PARP signals are shown, as indicated. (B) FACS analysis of IMR-32 cells treated as in (A) and stained with PI. The region used to evaluate the fraction of cells displaying sub-G1 DNA content is indicated. (C) Morphological appearance of IMR-32 cultures by bright-field microscopy (20x magnification), treated as in (A). Scale bar: 100 μm.

Together, these results indicate that PHA-680626 is an effective candidate to prevent the AURKA/N-Myc interaction in cultured cells, triggers N-Myc degradation and impairs the viability of neuroblastoma cell lines that are N-Myc-addicted for their proliferation.

## DISCUSSION

Poor prognosis for neuroblastoma patients is often associated to elevated levels of N-Myc, a transcriptional factor that mainly acts in favor of cellular proliferation, metastasis, cell cycle progression and protein translation (Dang, 2012). In spite of the paramount importance of this oncoprotein, the very low druggability of the latter has so far prevented any successful strategy involving N-Myc as a potential target for chemotherapeutic intervention. However, recent studies demonstrating that the physical interaction between the mitotic kinase AURKA and N-Myc sequesters the latter from proteolytic degradation (Richards et al., 2016; Büchel et al., 2017) paved the way for a new and promising chemotherapeutic approach in neuroblastoma. These findings have prompted the identification of compounds able to disrupt the AURKA/N-Myc complex, in order to promote the degradation of the latter (Meyerowitz et al., 2015; Brockmann et al., 2015)(Gustafson et al., 2014). Already known competitive inhibitors of AURKA, targeting the kinase active site, were demonstrated to also inhibit such interaction. Among these, CD532, MLN8054 and Alisertib, the best characterized compounds so far, are able to alter the conformation of AURKA in a way that prevents N-Myc binding, and for this reason, they have been hailed as “conformation disrupting” (CD) or “amphosteric” inhibitors (Meyerowitz et al., 2015).

As the amphosteric effects are neglected in most current inhibitor screenings (Gustafson et al., 2014), we investigated already known AURKA inhibitors with the aim of 1) expanding the repertoire of AURKA/N-Myc amphosteric compounds, and 2) at the same time, providing additional insights for their observed chemotherapeutic effects. To this end, we started this study from structural data mining on PDB, to identify crystal structures of AURKA whose conformation, due to the complex with active site inhibitors, was unsuited for N-Myc binding. The comparison of such AURKA structures with apo-AURKA (PDB: 4J8N), showed that the N- and C-terminal lobes of the kinase domain can adopt a very diverse set of orientations, ranging from ~85° in the case of MLN8054, to ~93° for CD532, and different conformations of the A-loop in the closed state, confirming that the view of kinases existing mainly in two states, “active” or “inactive”, based on the DFG-motif orientation (DFG-in -> active, DFG-out -> inactive) is somewhat simplistic. For example CD532, the best characterized amphosteric inhibitor so far, and the one exerting at the same time the amplest opening of the AURKA lobes and the most dramatic effects on N-Myc degradation, is paradoxically also a “DFG-in” inhibitor, according to the position of the DFG motif (Meyerowitz et al., 2015).

Those compounds that were predicted to disrupt the AURKA conformation, in a way similar to MLN8054/CD532, were obtained and further characterized. A series of initial kinase activity assays confirmed the inhibitory ability of each compound against AURKA, with IC_50_ in close proximity to the ones reported in literature with different *in vitro*/*in vivo* assays (Martin et al., 2012; Gustafson et al., 2014; Ferguson et al., 2017). MLN8054 (highly similar to Alisertib), with a 1-digit nanomolar IC_50_ value, was the most potent AURKA inhibitor tested, and it was adopted as archetypal amphosteric inhibitor in the next comparative analyses.

Using SPR, a well-established technique that yields a straightforward characterization of protein-protein and protein-ligand interactions, we confirmed *in vitro* that the two compounds identified by structural analysis as potential amphosteric inhibitors, are indeed able to interfere to with AURKA/N-Myc formation. In one case (i.e., Alisertib), where it was still possible to measure an apparent Kd_N-Myc_, the observed effect was to lower the affinity of N-Myc for the kinase. All the other investigated compounds were able to completely inhibit the interaction between the two proteins at saturating concentrations (10 times higher than their Kd). PHA-680626 was responsible for a complete dissociation of the complex. A closer inspection of the crystal structure and a detailed comparison of PHA-680626 with the highly similar PHA-739358 (Danusertib), the structure of which was solved in complex with AURKA (PDB 2J50), provided a rationale to the different behavior of the two closely related compounds (**Supplementary Figure 2A**). Indeed, the thiophene moiety of PHA-680626 is able to stabilize the A-loop in a close conformation that is unproductive for N-Myc binding, thanks to a stacking interaction with His280 of the A-loop. Conversely, the bulkier methoxy moiety of Danusertib hampers such interaction and prevents the A-loop of AURKA to adopt a “closed” conformation. Notably, topologically similar hydrophobic interactions were also observed in the structure of AURKA/MLN8054 and predicted via modeling (**Supplementary Figure 2B**) in the case of the fluorophenyl moiety of Alisertib (residues Val147, Ser278 and Val279), thus suggesting such interaction as an important pharmacophore feature for stabilizing the closed conformation of the A-loop and conferring an amphosteric behavior to the inhibitor. It is interesting to note that CD532 achieves a similar conformational change of the A-loop, without engaging any stabilizing contact with the latter. Indeed, the deeply buried 3-trifluoromethyl-biphenyl urea moiety of CD532 induces this inactive conformation via displacement and interactions with the β1 and β2 strands of the N-terminal lobe of AURKA, without reorienting the DFG motif, but wide-opening in this way the N-terminal lobe of the kinase, creating a large cleft in which the A-loop is stabilized in an inactive orientation through a unique network of hydrogen bonds. Therefore, taken together with structural data, these observations suggest that the conformational disruption of AURKA that is necessary for dissociate N-Myc can be achieved by adopting two complementary, but distinct strategies: stabilizing the A-loop in a closed conformation via formation of direct contacts, and/or wide-opening the N-terminal lobe of the kinase to create an ideal cleft for A-loop closure.

It is important to note that, in any case, fairly high concentrations of inhibitors were necessary to hamper the formation of the AURKA/N-Myc complex. Such high concentrations, which are necessary to unveil the amphosteric potential of AURKA inhibitors, suggest the difficulty that small inhibitors encounter when competing with the AIR of N-Myc. It was previously shown that AURKA is in dynamic equilibrium between active and inactive A-loop conformations and that the position of this equilibrium can be shifted by the binding of other ligands (Gilburt et al., 2017). The kinetics of N-Myc binding to AURKA (**Figure 2F**) measured by means of SPR suggests that an initial low association rate (K_on_) is then followed by a quite stable complex that shifts the dynamic equilibrium of AURKA in a conformation disfavoring successive binding of CD inhibitors. In this scenario, our data suggest that an amphosteric inhibitor of AURKA with a very low dissociation rate (in our study PHA-680626, for example, is the one with the lowest K_off_), which is independent on the concentration of the compound, might be more effective at displacing N-Myc. Recently, it has been shown that covalent Coenzyme A modification of AURKA via binding to the Cys290 residue of the A-loop is specific, and provides the proof of concept for a potential “dual anchor” irreversible inhibitory mechanism of AURKA (Tsuchiya et al., 2020). Such suicide inhibition would show the lowest reachable K_off_ value and would possibly keep the A-loop of AURKA in a conformation unsuitable for N-Myc binding, relieving from the need of high inhibitor concentrations.

Most importantly, our study expanded the repertoire of AURKA/N-Myc amphosteric compounds and found that two other compounds other than CD532, Alisertib and MLN8054, i.e. PHA-680626 and RPM1722, already known as orthosteric inhibitors, are indeed able to prevent N-Myc binding to AURKA. When moving to cellular studies, PHA-680626 proved effective both in inducing a cell cycle arrest comparable to MLN8054 and in destabilizing N-Myc. This was observed in a N-Myc inducible U2OS cell line and in neuroblastoma IMR32 cells, where cytotoxic effects were also observed. Surprisingly, no effect was induced by RPM1722 treatment, including no observable changes in the cell cycle phases distribution, which would be expected for an AURKA inhibitor; it is therefore possible that this compound does not efficiently enter the cell. To the best of our knowledge, no cell permeability studies were carried out so far on RPM1722. It is conceivable therefore that further optimization of this scaffold will be required, involving the improvement of cell permeability, by rational-based modifications of the original scaffold (e.g., carrier-linked prodrugs). On the other hand, the promising behavior of PHA-680626 suggests that its use in the treatment of neuroblastoma can be further explored. As discussed above, although the structural analysis of the highly similar Danusertib suggests that the latter is unable to exert the same effects of PHA-680626, nevertheless it will be interesting to investigate in future studies whether such “amphostericity” is a common attribute of this class of compounds. The recent report of successful combined use of AURKA CD-inhibitors and ATR inhibitors in MYCN amplified neuroblastoma (Roeschert et al., 2021), based on the S-phase function of the AURKA/N-Myc complex, strengthen the potential value of using compounds disrupting both AURKA activity and the interaction with N-Myc for the treatment of this aggressive tumor type.

## MATERIALS AND METHODS

### Screening on Protein Data Bank and Homology Modeling

CD532 (PDB entry:4J8M) and MLN8054 (PDB entry: 2×81) were used as prototype of amphosteric inhibitors to quantitatively measure the magnitude of the conformational disrupting effect, as suggested in Meyerowitz et al., 2015. The opening between the N-terminal domain, compared to the C-terminal lobe, was asserted by measuring the dihedral angle between the α-carbons of V324, E308, and A172, comparing it to the one measured for apo-AURKA (empty binding pocket, 4J8N, angle: 81.7°) (Gustafson et al., 2014). In order to generate the ensemble of crystallographic structures for our analysis, we have conducted a data mining of all the available entries of AURKA kinase on Protein Data Bank (PDB). We have identified 198 structures for AURKA kinase overall and, in 95 of them, the kinase in complex with already known orthosteric inhibitors (March, 2018). Subsequently, we have conducted a visual examination of the identified structures using the open-source molecular visualization system PyMOL (The PyMOL Molecular Graphics System, Version 1.2r3pre, Schrödinger, LLC).

We proceeded aligning every protein-inhibitor complex with the PDB entries 4J8M and 2×81 in order to find other molecules that can both open the “angle” of Aurora-A and induce a complete flip of the activation loop, retaining it in a “closed” conformation not able to bind N-Myc. We have excluded from our examination all the structures that lacked resolved residues at the key N-Myc binding regions; proceeding this way we have selected 10 entries (PDB codes: 2J4Z, 2J50, 2×6E, 2×81, 2WTV, 3UOJ, 3UOH, 3UNZ, 3UO6, 4J8M, 5ONE). We have then conducted another screening to exclude the “doublet” entries, in which Aurora-A is in complex with the same molecule (e.g., 2×81 and 2WTV in complex with MLN8054) and to consider only the best candidate in a family of inhibitors with the same backbone and the same structural change (e.g., 3UO6, 3UOJ, 3UOH, 3UNZ have the same chemical scaffold, but RPM1722 (3UOH) has the best affinity for AURKA, with a K_d_=13 ± 2.2 nM). The final dataset was formed by four structures: 4J8M (Gustafson et al., 2014), 2×81 (Sloane et al., 2010), 2J4Z (Fancelli et al., 2006), 3UOH (Martin et al., 2012). The crystal structure of AURKA in complex with MLN8054 (PDB: 2×81) was used as a starting point to generate the model of AURKA in complex with Alisertib, using the “homology modeling” tool of PyMod 3.0 (Janson & Paiardini, 2020).

### Purification of Aurora-A Catalytic Domain

For the purification of the catalytic domain of human AURKA (amino acids 122-430), the plasmid pETM11 was used. The resulting protein contains an N-terminal His tag and an intervening TEV protease cleavage site. For the pre-inoculation, more colonies were taken from a plate and resuspended in LB containing 40μg/ml kanamycin, with growth at 37°C overnight. After inoculation 1:100 in 2L of LB, culture was grown at 37°C for 4 hours and 30 minutes. When OD_600nm_ achieved the value of 0.5, the expression of the recombinant protein was induced with 0.2 mM IPTG and the culture was incubated at 28°C overnight. The pellet, after being centrifuged at 5500 rpm, at 4°C for 20 min, was resuspended in about 130 ml of 50 mM Hepes, pH 7.4, 5 mM MgCl_2_, 300 mM NaCl, 10% glycerol (buffer A) and 1 cOmplete Protease Inhibitor Cocktail tablet (SIGMA-Aldrich), sonicated in ice (for 2 min, 20 sec pulse on and 20 sec pulse off) and centrifuged at 12000 rpm for 25 min at 4°C, twice. The obtained supernatant was filtered and loaded onto a 5-ml HisTrap column (GE Healthcare), previously equilibrated with buffer A (flow rate 1ml/min). The column was washed with 10 ml buffer A and then with buffer A containing 20 mM, 40 mM and 100 mM imidazole. Elution was performed with buffer A + 300 mM imidazole. Fractions containing AURKA were collected and dialyzed against Buffer A, at 4°C. In order to use the kinase for the activity assays, it was essential to phosphorylate the purified catalytic domain, so to activate it. That was accomplished by incubation of AURKA with 400μM of ATP (>10fold of K_m_) for 3 hours on ice. To remove the ATP in excess, the protein was dialyzed in Buffer A overnight at 4°C. Once activated, AURKA was aliquoted and stored at −80°C, without losing stability or activity.

### Chemistry

Compounds for testing were purchased from Molport (https://www.molport.com/). All chemicals were of the highest purity available (guaranteed purity of over 90% by H-NMR or LCMS), as certified by vendor. PHA-680626 was a gift by Nerviano Medical Sciences Srl. The synthesis of RPM1722 (RDS4007) was carried out as reported in Scheme 1 (see Supplementary Information), using the 2,4-dichloropyrimidine as starting material. Two subsequent aromatic nucleophilic substitutions were performed. First, the 2-bromoaniline was introduced in position 4 of the pyrimidine ring in the presence of aqueous hydrochloric acid (1.0 M), allowing the reaction to proceed for 36 hours at room temperature. Then, the second chlorine atom was substituted with 4-aminobenzoic acid, by means of a microwave-assisted reaction performed using absolute ethanol as a solvent at 100°C for 20 minutes. The synthesis was carried out providing some changes to a previously published work (Lawrence et al., 2012). The synthetic scheme, the procedures and the analytical spectroscopic data are reported in the Supplementary Information.

### Kinase Activity Assays

All kinase assays for AURKA catalytic domain were performed with ADP-Glo™ Kinase Assay Kit (Promega©). For Alisertib the concentrations tested were: 0.075nM, 0.155nM, 0.312nM, 0.665nM, 1.250nM, 2.5nM, 5nM, 10nM, 20nM and 40nM; for CD532: 3.12nM, 6.25nM, 12.5nM, 25nM, 50nM, 100nM, 200nM, 400nM, 800nM and 1600nM; for MLN8054: 0.15, 0.31, 0.62nM, 1.25nM, 2.5nM, 5 nM, 10nM, 20nM, 40nM, 80nM and 160nM, for PHA680626: 3.12nM, 6.25nM, 12.5nM, 25nM, 50nM, 100nM, 200nM, 400nM, 800nM and 1600nM; for RMP1722 were: 0.5nM, 1nM, 2nM, 4nM, 8nM, 16nM, 32nM and 64nM. The reaction mixture (20 μl) contained the different concentrations of each inhibitor (obtained by adding 1μl of inhibitor solubilized in 100% DMSO), 35 μM ATP, 100 ng AURKA in 16 mM Tris HCl, pH 7.5, 8 mM MgCl_2_, 0.04 mg/ml BSA, 0.2 mM DTT. After incubation at room temperature for 30 min, the reaction was started by adding 5 μl of Myelin Basic Protein (20 mg/ml), a substrate of AURKA and incubated at 30°C. Then, the assay was performed in two steps; first, an equal volume (25 μl) of ADP-Glo™ Reagent was added to terminate the kinase reaction and deplete the remaining ATP. Second, the Kinase Detection Reagent (50 μl) was added to simultaneously convert ADP to ATP and allow the newly synthesized ATP to be measured using a luciferase/luciferin reaction. The luminescence produced at the end of every reaction, which corresponds to the kinase activity of AURKA, was measured as relative light unit (RLU) with a Luminoskan™ Microplate Luminometer (Thermo Scientific). Data analysis was carried out with the PRISM8 (Graph Pad) software, using the following equation:

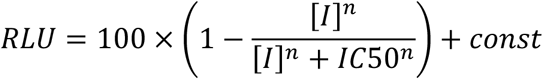

from what we obtained the IC50, with [I] as the concentration of inhibitor, *n* the Hill coefficient, and const as residual activity.

### Surface Plasmon Resonance

SPR experiments were performed at 25 °C using a Pioneer AE optical biosensor (Molecular Devices-ForteBio) equipped with a PCH sensor chip and equilibrated with running buffer 10mM Hepes pH7.4, 150 mM NaCl, 1% DMSO, 0.005% Tween20. Sensor chip was chemically activated for 7 min by injecting 175 μL of a 1:1 mixture of 100 mM N-hydroxysuccinimide (NHS) and 400 mM ethyl-3(3-dimethylamino) propyl carbodiimide (EDC) at a flow rate of 25 μl/min. AURKA was immobilized on both Ch1 and Ch3 channels of the activated sensor chip by using standard amine-coupling procedures (Lundström, 1994). Flow cell Ch2 was left empty and used as control. Briefly, a 0.1 mg/mL AURKA solution (in 10 mM sodium acetate, pH 4.5) were injected at 10μL/min on channels 1 and 3 (channel 2 was used as reference), followed by a 70-μL injection of 1M ethanolamine pH 8.0 to block any remaining activated groups on the surface. AURKA was captured to a density of ~14000 resonance units (RUs) on both Ch1 and Ch3 flow cells. The stability of the AURKA surface was demonstrated by the flat baseline achieved at the beginning (0–60s) of each sensorgram. To correct for bulk refractive index shifts, a DMSO calibration plot was also constructed (buffer sample containing 0.5–1.5% DMSO) (Frostell-Karlsson et al., 2000).

Compounds for testing were solubilized in 100% DMSO and then diluted in 10 mM Hepes pH 7.4, 150 mM sodium chloride and 0.005% Tween 20 up to a final concentration of 1% DMSO (running buffer) or 4% (CD532) and injected at different concentrations onto the sensor chip at a constant flow rate of 150 μl/min. A dissociation of 180 sec was used and a mild regeneration of the surfaces was carried out every three injections by using a solution of 1M NaCl and 10mM NaOH. At least two experiments for each analyte were performed. The interaction of the tested inhibitors with immobilized AURKA was detected as a measure of the change in mass concentration, expressed in RUs. The same conditions were used for the competition experiments between the compounds and N-Myc, respectively. In this case, N-Myc was diluted in running buffer containing the inhibitor at a constant concentration 10 times higher than its Kd. Different concentrations of N-Myc (ranging from 125 nM to 16 μM) were then injected on the sensor chip at a constant flow of 150 μl/min of running buffer containing the inhibitor at saturating concentration. All sensorgrams were processed by using double referencing (Myszka & Morton, 1998). First, responses from the reference surface (Ch2) were subtracted from the binding responses collected over the reaction surfaces to correct for bulk refractive index changes between flow buffer and analyte sample. Second, the response from an average of the blank injections (zero analyte concentrations) was subtracted to compensate for drift and small differences between the active channel and the reference flow cell Ch2 (Myszka, 1999). To obtain kinetic rate constants and affinity constants, the corrected response data were fit in the QDAT software. A kinetic analysis of each ligand/analyte interaction was obtained by fitting the response data to a reversible 1:1 bimolecular interaction model. The equilibrium dissociation constant (Kd) was determined by the ratio k_off_/k_on_.

### Cell lines, synchronization protocols and treatments

Cell lines were grown at 37°C and 5% CO_2_. For the generation of the human U2OS/MYCN-tFT osteosarcoma cell line, N-Myc (Addgene #74163) was cloned into plasmid pcDNA5/FRT/TO (Thermo Fisher Scientific) bearing an mCherry and mNeon tandem fluorescent tag (tFT, Khmelinskii et al. 2012; kind gift of Michael Knop, ZMBH, University of Heidelberg). This construct was used to create the N-Myc-tFT cell line by Flp recombinase-mediated integration into the genome of a U2OS FRT/TO Flp-In host cell line (kind gift of Adrian Saurin, Dundee) according to the manufacturer’s instructions for the Flp-InTM T-RExTM System (Thermo Fisher Scientific). This cell line was grown in complete DMEM (Dulbecco’s Modified Eagle Medium) supplemented with 10% tetracycline-free fetal bovine serum (FBS) and the induction of N-Myc expression was obtained by adding 1 µg/ml doxycycline (tetracycline analogue; Santa Cruz Biotechnology). Treatment with 9 µM Ro-3306 (Sigma-Aldrich, SML0569) for 22 h or with 10 µM MG132 (Chem-Cruz, sc-201270) for 4 h was performed when indicated. The IMR32 neuroblastoma cell line (kind gift of Prof. Giuseppe Giannini, Sapienza University of Rome) was grown in MEM (Minimum Essential Medium) supplemented with 10% FBS and 1% of non-essential amino acids. Both cell lines were treated with 1 µM of inhibitors (MLN8054, PHA-680626, RPM1722), or 0,1% DMSO as control, for 22 h or 48 h, as indicated. Images of IMR32 cultures after treatments were acquired using a ZOE fluorescent cell imager (Biorad).

### *In Situ* Proximity Ligation Assays (*is*PLA)

*In situ* proximity ligation assays (*is*PLA) were performed on cells grown on coverslips and fixed with 3.7% formaldehyde/30 mM sucrose in PBS, 10 min at room temperature, followed by permeabilization in PBS containing 0.1% TritonX-100, 5 min at room temperature. The Duolink PLA kit (DUO92007, Sigma-Aldrich) was used according to manufacturer’s instructions. The amplification time was 100 minutes and the primary antibodies pair to detect the interaction was rabbit anti-Aurora-A (14475, Cell Signaling, 1:100 o.n.) and mouse anti-N-Myc (NCM II 100, Calbiochem, 1:20). Counterstaining. with DAPI was performed at the end of the procedure. For quantification of *is*PLA fluorescence signals, samples were analyzed with an inverted microscope Eclipse Ti (Nikon) using a Clara camera (ANDOR technology) and 60x (oil immersion, N.A. 1.4) objective along the z-axis every 0.4 μm for a range of 3.2 μm. Images were acquired in automated mode with the JOBS module of the Nis-Elements H.C. 5.11 software. The “general analysis” module of Nis Elements H.C. 5.11 was then used for automatic spot count; *is*PLA signals were identified based on fixed parameters in all images and those within nuclei (defined by DAPI signal) were counted. Elaboration of the images was made by Adobe Photoshop CS 8.0 and statistical analyses were performed with GraphPad; specific tests are indicated in Figure legends.

### Western Blotting

For Western Blot analysis, synchronized or asynchronously growing cells were lysed in RIPA buffer (50 mMTris-HCl pH 8.0, 150 mM NaCl, 1% NP40, 1 mM EGTA, 1 mM EDTA, 0.25% sodium deoxycholate) supplemented with protease and phosphatase inhibitors (Roche Diagnostic). Proteins were resolved by 10% SDS PAGE and transferred on a nitrocellulose membrane (Protran BA83, GE Healthcare) using a semi-dry system (Bio-Rad Laboratories S.r.l.). About 30 µg of extract per lane was loaded. Antibodies were: mouse anti-N-Myc (NCM II 100, Calbiochem, 1:100); rabbit anti-Lamin B1 (ab16048, Abcam, 1:4000); rabbit anti-PARP (9542, Cell Signaling, 1:1000 o.n.). HRP-conjugated secondary antibodies (BioRad) were revealed using the Clarity Western ECL substrate (BioRad, 170-5061).

### FACS analysis

To analyze cell-cycle phases distribution, both non-adherent and adherent cells were collected by centrifugation, washed with PBS and fixed with 50% methanol overnight at +4°C. Cell cycle phase distribution was analyzed after incubation for 30 minutes in the dark with propidium iodide (PI, 0.03 mg/ml) and RNAsi A (0.2 mg/ml) using a flow cytofluorimeter Epics XL apparatus (Beckman Coulter). Cell aggregates were gated out on bi-parametric graph FL-3lin/ratio as described in (Ferrara et al., 2018). Cell samples were analyzed in a Coulter Epics XL cytofluorometer (Beckman Coulter) equipped with EXPO 32 ADC software. At least 10 000 cells per sample were acquired. The percentage of cells in the different phases of cell-cycle and in sub-G1 compartment was calculated using Flowing Software 2.5.1.

## Acknowledgments

We thank I.A. Asteriti for experimental support and insightful discussions. The authors wish to thank Richard Bayliss and Dr. Mark Richards for assisting this study by providing the vector pETM6T1, coding for the 28-89 peptide of N-Myc, the vector pETM11 coding for Aurora-A Catalytic Domain (aa 122-403) as well as for their interesting advice, discussions and precious help. Thanks to Michael Knop for tFT plasmids and to Adrian Saurin for U2OS FRT/TO cells. This work is dedicated to the memory of our beloved mentors Prof. Francesco Bossa and Prof. Donatella Barra.

## Funding Statement

AP received support from Associazione Italiana Ricerca sul Cancro (AIRC, https://www.airc.it/) MFAG 20447 and Progetti Ateneo Sapienza University of Rome (https://www.uniroma1.it). AP acknowledges the CINECA award under the ISCRA initiative, for the availability of high-performance computing resources and support (IsC68_altmod). The funders had no role in study design, data collection and analysis, decision to publish, or preparation of the manuscript.

## Notes

### Competing Interest Statement

The authors have declared no competing interest.

### Summary of Updates

Figure 4 revised

