## Supplementary Figures for "A Focused Small-Molecule Screen Identifies PHA-680626 as an Amphosteric Inhibitor Disrupting the Interaction between Aurora-A and N-Myc"

Supplementary Figure 1

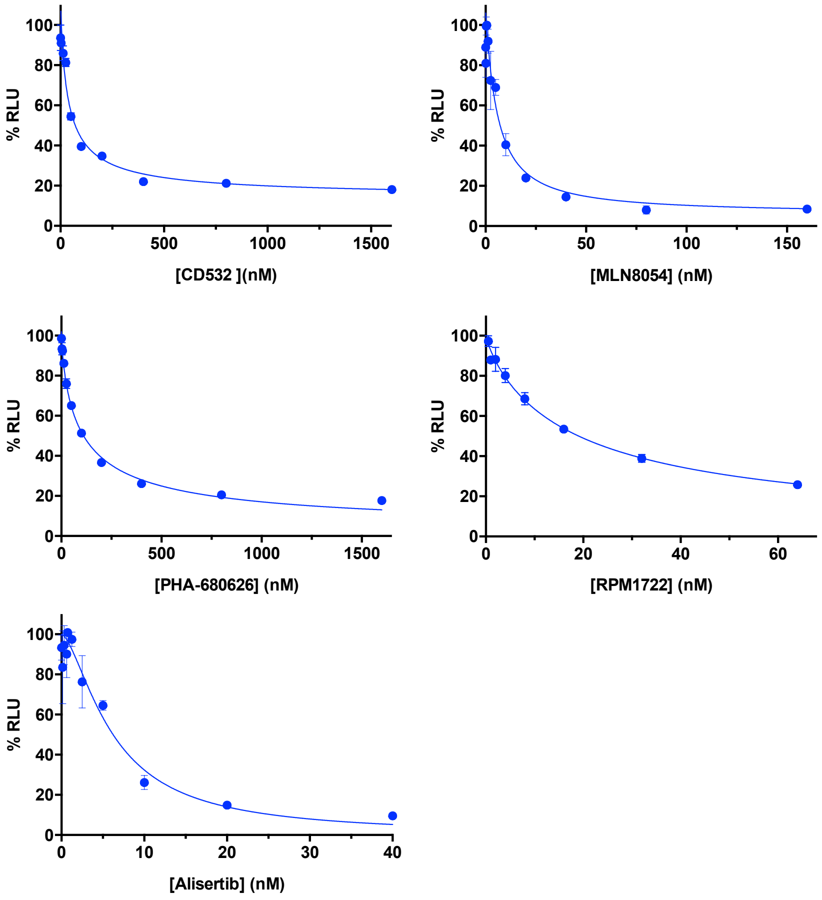


Supplementary Figure 1. Inhibition assay of tested inhibitors against AURKA. The activity of phosphorylated AURKA, expressed as relative light units (RLU) and normalized to 100%, was measured in the presence of increasing concentrations of inhibitors.

Supplementary Figure 2

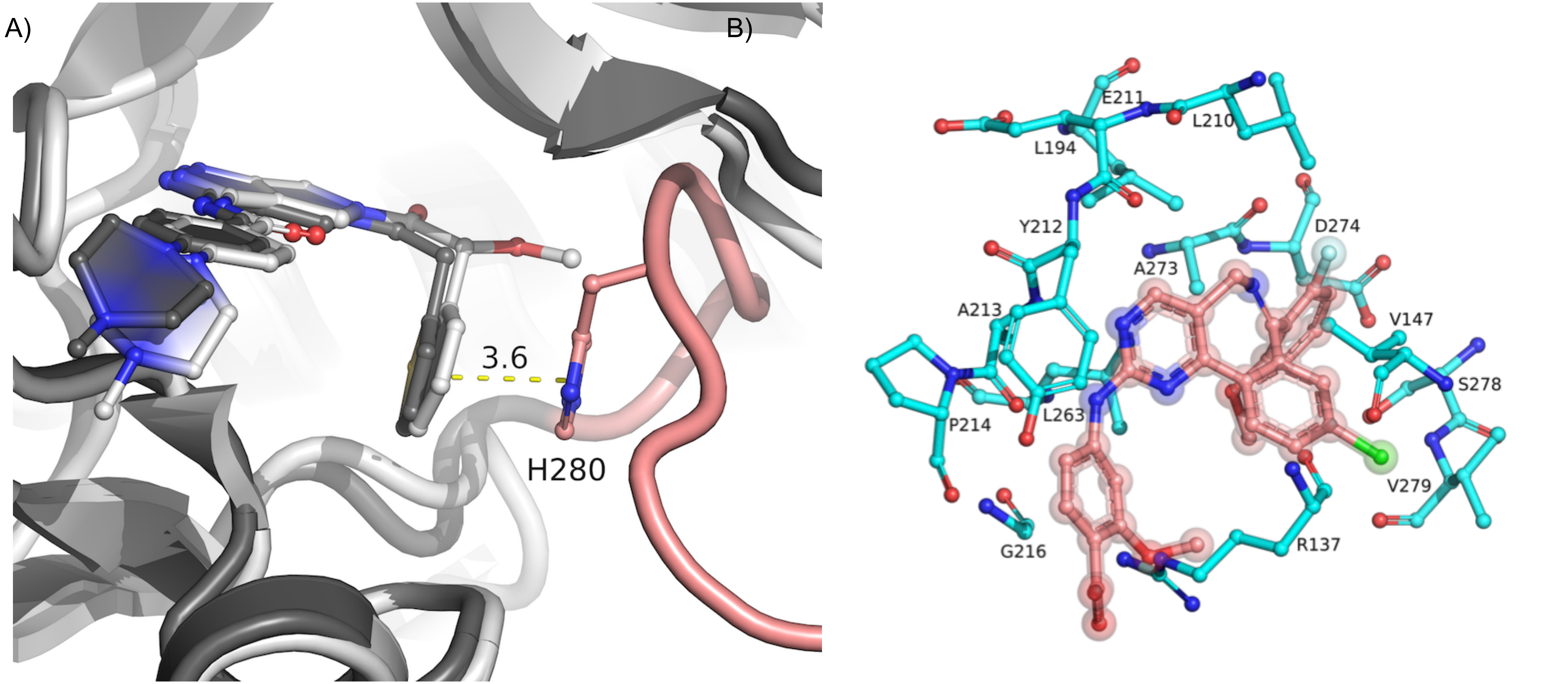


**Supplementary Figure 2.** A) Structural comparison of PHA-739358 (Danusertib) and PHA-680626. The thiophene moiety of PHA-680626 (dark grey sticks) is able to stabilize the A-loop (in pink) in a closed conformation that is unproductive for N-Myc binding, thanks to a stacking interaction (dashed yellow line labeled with distance in Å) with His280. In contrast, methoxy moiety of PHA-739358 (dark grey sticks) hampers such interaction and prevents the A-loop of AURKA to adopt a close conformation. B) Modeling of Alisertib (pink sticks) into the active site of human AURKA (cyan sticks). Alisertib is predicted to form hydrophobic interactions with residues Val147, Ser278 and Val279, and stabilize the A-loop in a closed conformation.

Supplementary Figure 3

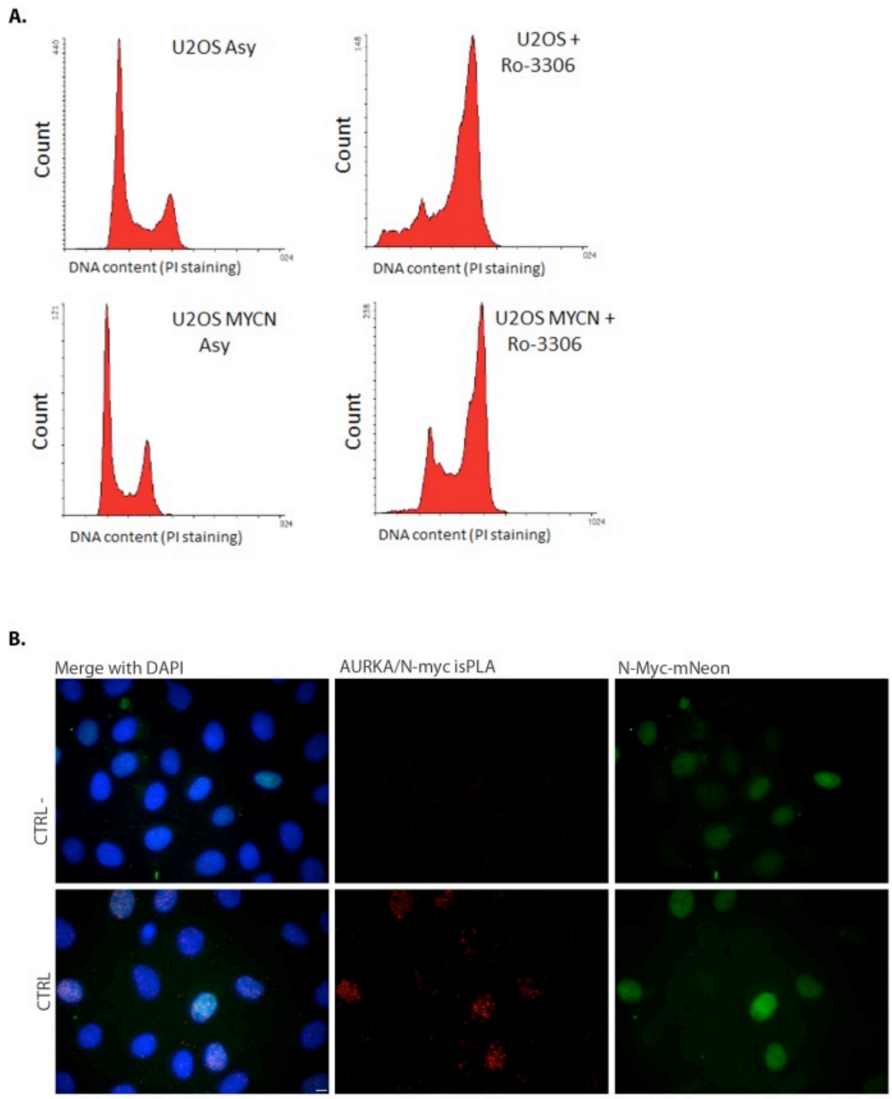


**Supplementary Figure 3**. **Control conditions for experiments shown in Figure 4.** (A) FACS analysis of U2OS and U2OS MYCN cell lines (propidium iodide staining) following treatment with RO-3306 for 22 h. (B) Representative images of *is*PLA in the U2OS MYCN cell line, under standard conditions (CTRL) or carried out without the AURKA primary antibody (CTRL-).

Supplementary Scheme 1. Synthesis of RPM1722

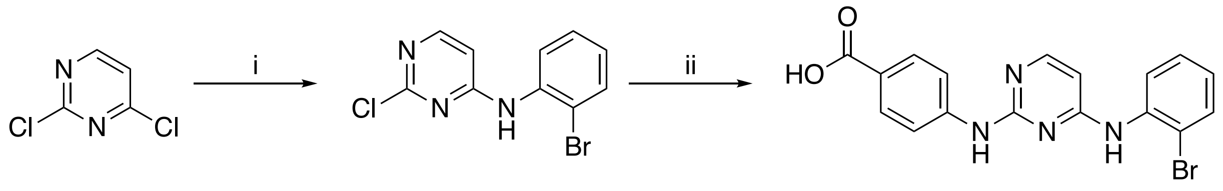


[a]Reagents and conditions: (i) 2-bromoaniline, HCl 0.1M aq, room temp, 36 h, 80.4% yield; (ii) 4-aminobenzoic acid, $EtOH_{(abs)}$, 100 °C, 20 min, microwave, 72.5% yield.

**Chemistry: General**. Melting points were determined on a Bobby Stuart Scientific SMP1 melting point apparatus and are uncorrected. IR spectra were recorded on a PerkinElmer Spectrum-One spectrophotometer. 1H nuclear magnetic resonance (NMR) spectra were recorded at 400 MHz on a Bruker AC 400 Ultrashield 10 spectrophotometer (400 MHz). Dimethyl sulfoxide-*d6* 99.9% (CAS 2206-27-1) of isotopic purity (Aldrich) was used. Mass spectra were recorded on a ThermoFinnigan LCQ Classic liquid chromatography−tandem mass spectrometry (LC−MS/MS) ion trap equipped with an electrospray ionization (ESI) source and a syringe pump. Samples ($10^{-4}$–$10^{-5}$ M in methanol/water 9:1) were infused in the electrospray system at a flow rate of 5−10 μL $min^{-1}$. Column chromatographies were performed on silica gel (Merck; 70−230 mesh). All compounds were routinely checked with thin-layer chromatography by using aluminum-baked silica gel plates (Merk silica gel plates on aluminium support 60 F254). Developed plates were visualized by UV light. Solvents were reagent grade and, when necessary, were purified and dried by standard methods. Concentration of solutions after reactions and extractions involved the use of rotary evaporator (Büchi) operating at a reduced pressure (ca. 20 Torr). Organic solutions were dried over anhydrous sodium sulfate (Merck). All solvents were freshly distilled under nitrogen and stored over molecular sieves for at least 3 h prior to use. Analytical results agreed to within ±0.40% of the theoretical values. Microwave reactions were conducted using a CEM Discover system unit (CEM. Corp., Matthews, NC). The machine consists of a continuous focused microwave-power delivery system with operator selectable power output from 0 to 300 W. The temperature of the contents of the vessel was monitored using a calibrated infrared temperature control mounted under the reaction vessel. All experiments were performed using a stirring option whereby the contents of the vessel are stirred by means of a rotating magnetic plate located below the floor of the microwave cavity and a Teflon-coated magnetic stir bar in the vessel.

*Synthesis of N-(2-bromophenyl)-2-chloropyrimidin-4-amine* (**RDS 4005**). To a suspension of 2,4-dichloropyrimidine (1.00 g, 6.71 mmol) in an aqueous solution of HCl 0.1 M (15 mL), 2-

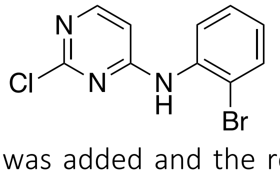

bromoaniline (1.13 g, 6.58 mmol) was added and the reaction was allowed to stir at room temperature for 36 hours. Upon completion of the reaction, the acidic suspension was neutralized with saturated solution of $NaHCO_3$ and extracted with ethyl acetate (2x150 mL). The organic layer was washed with saturated solution of sodium chloride (2x150mL), dried

over sodium sulphate and evaporated under reduced pressure to afford the crude product which was purified by silica gel chromatography n-hexane/ethyl acetate (3:2) to afford the title compound (1.50g, yield: 80.4%) as an off-white solid (m.p. 137 °C). $^{1}$H NMR (400 MHz, DMSO-$d_6$) $\delta$ 9.73 (bs, 1H, NH), 8.13 (d, $J$ = 5.8 Hz, 1H, pyrimidine), 7.73 (dd, $J$ = 1.5 Hz, 7.9 Hz, 1H, Ar), 7.53 (dd, $J$ = 1.5 Hz, 7.9 Hz, 1H, Ar), 7.44 (td, $J$ = 1.5 Hz, 7.9 Hz, 1H, Ar), 7.23 (td, $J$ = 1.5 Hz, 7.9 Hz, 1H, Ar), 6.59 (d, $J$ = 5.8 Hz, 1H, pyrimidine); IR $\nu$ 3186 (NH) cm$^{-1}$; MS: *m/z* (ESI) calcd for $[C_{10}H_7BrClN]^+$: 284.54, found, 284.05. Anal. Calcd for $C_{10}H_7BrClN_3$: C, 42.21; H, 2.48; Br, 28.08; Cl, 12.46; N, 14.77%. Found: C, 42.23; H, 2.49; Br, 28.06; Cl, 12.47; N, 14.75%.
SMILES: ClC1=NC=CC(NC2=C(Br)C=CC=C2)=N1

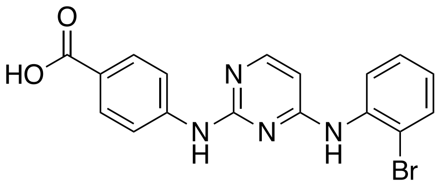


Synthesis of *4-((4-((2-bromophenyl)amino)pyrimidin-2-yl)amino)benzoic acid* (**RDS 4007**; **RPM1722**). A mixture of chloropyrimidine (**RDS 4005**) (0.100g, 0.350 mmol) and 4-aminobenzoic acid (0.048g, 0.350 mmol) in absolute ethanol (0.5 mL) was heated in a microwave reactor at 100 °C for 20 minutes. The resulting white precipitate was filtered and purified by silica gel chromatography chloroform/methanol (8.5:1.5) to afford the title compound (0.098g, yield: 72.5%) as a white solid (m.p. 256-257 °C). $^{1}$H NMR (400 MHz, DMSO-$d_6$) $\delta$ 12.52 (bs, 1H, COOH), 9.59 (s, 1H, NH), 9.12 (s, 1H, NH), 8.13 (d, $J$ = 6.5 Hz, 1H, pyrimidine), 7.77-7.81 (m, 6H, Ar), 7.52 (td, $J$ = 1.6 Hz, 7.8 Hz, 1H, Ar), 7.29 (td, $J$ = 1.5 Hz, 7.9 Hz, 1H, Ar), 6.36 (d, $J$ = 6.5 Hz, 1H, pyrimidine); IR $\nu$ 3405 (NH), 2437 (COOH), 1600 (C=O carboxylic acid) cm$^{-1}$; MS: *m/z* (ESI) calcd for $[C_{17}H_{13}BrN_4O_2]^+$: 385.22, found, 385.50. Anal. Calcd for C, 53.01; H, 3.40; Br, 20.74; N, 14.54; O, 8.31%. Found: C, 53.09; H, 3.41; Br, 20.70; N, 14.52%.
SMILES: OC(C(C=C1)=CC=C1NC2=NC=CC(NC3=C(Br)C=CC=C3)=N2)=O
